# Assessing spontaneous sensory neuron activity using *in vivo* calcium imaging

**DOI:** 10.1101/2023.01.12.523728

**Authors:** Sonia L Ingram, Kim I Chisholm, Feng Wang, Yves De Koninck, Franziska Denk, George L Goodwin

## Abstract

Heightened spontaneous activity in sensory neurons is often reported in individuals living with chronic pain. It is possible to study this activity in rodents using electrophysiology, but these experiments require great skill and can be prone to bias. Here, we have examined whether *in vivo* calcium imaging with GCaMP6s can be used as an alternative approach. We show that spontaneously active calcium transients can be visualised in the fourth lumbar dorsal root ganglion (L4 DRG) via *in vivo* imaging in a mouse model of pain. Application of lidocaine to the nerve, between the inflamed site and the DRG, silenced spontaneous firing and revealed the true baseline level of calcium for spontaneously active neurons. We used this data to train a machine leaning algorithm to predict when a neuron is spontaneously active. We show that our algorithm is accurate in two different models of pain: intraplantar Complete Freund’s Adjuvant and antigen-induced arthritis, with accuracies of 90.0% +/-1.2 and 85.9 % +/-2.1, respectively, assessed against visual inspection by an experienced observer. The algorithm can also detect neuronal activity in imaging experiments generated in a different lab using a different microscope configuration (Accuracy = 94.0 % +/2.2). We provide a Google Colaboratory Notebook to allow anyone easy access to this novel tool, for assessment of peripheral neuron activity in their own calcium imaging setups.

## Introduction

Spontaneous activity in the nervous system is defined as the generation of neural activity without any appreciable external task or stimulus. In peripheral sensory neurons, this type of activity is rarely found in healthy individuals^1,2^. Instead, it is prevalent in many people with chronic neuropathic^3-5^ and musculoskeletal^6,7^ conditions, with a report indicating that the levels of spontaneous activity can directly correlate with levels of pain^8^. Consistent with these observations in humans, an increase in the proportion of sensory neurons with spontaneous activity has also been reported in rodent models of chronic pain^3,9-11^. Unsurprisingly, given this evidence, pharmaceutical companies have focused many efforts on developing analgesics that can normalise this hyperactivity, by targeting ion channels and receptors located in peripheral sensory neurons, in the hope of blocking the generation of unwanted pain^12,13^. And yet, this strategy has so far failed to produce novel analgesic medications. One possible reason why it might be difficult to make progress in this area, is that we lack basic mechanistic knowledge.

Traditionally, spontaneous activity has been studied using *in vivo* electrophysiology, which typically involves one-by-one recordings from individual sensory nerve filaments using a highly technically challenging teased-fibre setup. We are ourselves one of the few groups that are proficient in this technique^14,15^ and are therefore familiar with its laborious nature. Nociceptive C-fibres are particularly difficult to record from, due to their biophysical properties. Specifically, their small size and large resistance cause their extracellular spike amplitudes to be very small, and this requires the nerve to be teased into extremely fine filaments (∼10 µm diameter) to achieve a sufficiently high signal to noise ratio. Such technical challenges mean that studies of this kind are lengthy and low throughput, usually reporting only on around 10-15 fibres per animal. They are also prone to bias. For example, the smallest fibres are likely to be under-sampled, as they are the first to disintegrate when teased. Moreover, receptive field searching for nociceptors can cause neuronal sensitisation and an increase in the proportion of spontaneously active fibres found^16^.

*In vivo* calcium imaging is a relatively new method for studying sensory neuron function, which can be used to study hundreds of cells simultaneously^17^. Given the difficulties associated with studying spontaneous activity via electrophysiology, we set out to determine whether *in vivo* imaging could be used as an alternative, to improve our knowledge and understanding of this phenomenon. For this, we used the Complete Freund’s adjuvant (CFA) model of pain to observe spontaneously active calcium transients in mice whose sensory neurons were labelled with the calcium indictor GCaMP6s. To differentiate true spontaneous activity from unrelated fluctuations in calcium signals, we used the local anaesthetic lidocaine to block incoming electrical activity from the inflamed site. These data were then assessed by an experienced observer, to detect which neurons were spontaneously active (i.e. showed activity before, but not after lidocaine application). Using these “ground truth” data, we trained a machine learning algorithm to detect spontaneous activity in a standard recording protocol (without the need for lidocaine application). We confirmed the robustness of the algorithm by testing it on independent CFA data, an alternative pain model and on data generated using a different microscope setup in an independent laboratory. Our results suggest that *in vivo* imaging with GCaMP6s is suitable for large-scale assessment of spontaneous activity in sensory neurons.

## Materials and Methods

### Animals

Animal experiments conducted in the UK: Adult C57BL/6J male and female mice (n = 32; Charles River, UK) weighing 24-30 g were used in this study. Mice were housed on a 12/12 h light/dark cycle with a maximum of 5 mice per cage, with food and water available *ad libitum*. All experiments were performed in accordance with the United Kingdom Home Office Animals (Scientific Procedures) Act (1986). Animal experiments conducted in Canada: All experiments were performed in accordance with regulations of the Canadian Council and Animal Care. Homozygous MrgprD-Cre mice (B6.129S1(Cg)-*Mrgprd*^*tm1*.*1(cre)And*^/Mmnc, MMRRC, 036118-UNC) were crossed with C57BL/6J mice (JAX, 000664) to have MrgprD-Cre heterozygous mice. A total of 6 adult mice were used in 2-photon recording experiments.

### Administration of GCaMPs

We utilised the genetically encoded calcium indicator GCaMP6s for imaging sensory neuron activity^18^. GCaMP6s was delivered to sensory neurons via an adeno-associated viral vector of serotype 9 (AAV9), which was administered to mouse pups at P2-P7 as previously described^19^. Briefly, groups of 3-4 mice were separated from their mother. 5 µl of AAV9.CAG.GCaMP6s.WPRE.SV40 or AAV9.syn.GCaMP8s-WPRE virus (Addgene, USA) was injected subcutaneously in the nape of the neck, using a 10 µL Hamilton syringe with a 34G needle. Mice were separated from their mother after weaning and then used for *in vivo* imaging from 10 weeks after the injection. These mice were used for 1-photon imaging experiments.

For validation of our algorithm, we asked for existing data from another laboratory. These data had been generated for the purposes of an entirely different study. In that study, non-peptidergic nociceptors were specifically targeted for imaging, using intraplantar injections of 10 µl of AAV9.CAG.Flex.GCaMP6s.WPRE.SV40 (Addgene, 100842-AAV9) into newborn MrgprD-Cre+ mice. In contrast to our data, imaging was performed using a 2-photon imaging setup.

### CFA pain model

To induce acute inflammation and pain, 20 µL Complete Freund’s Adjuvant (CFA; 1 mg/ml, Sigma, UK) was injected intraplantar into the left hind paw using a 30G insulin syringe.

### Antigen Induced Arthritis model

Mice were immunised using an emulsion of CFA (3.3 mg/ml; Sigma, UK) and mBSA (40 mg/ml; Sigma, UK), as described previously^20^. Briefly, mice were anesthetised using 2 % isoflurane, and 100 µl of the emulsion was subcutaneously injected at the base of the tail and in the right flank (50 µl each). Mice were then allowed to recover and were returned to their home cages. Seven days after immunisation, mice were anaesthetised with isoflurane, and 2.5-5 µl of mBSA (200µg) was injected into the left knee joint using a Hamilton syringe with a 30G needle.

### *In vivo* imaging of sensory neuron activity using GCaMP6s

#### photon imaging

Mice were anesthetized using a combination of drugs: 1-1.25 g/kg 12.5% w/v urethane administered intraperitoneally and 0.5-1.5 % isoflurane delivered via a nose cone. Body temperature was maintained close to 37°C using a homeothermic heating mat with a rectal probe (FHC). An incision was made in the skin on the back, and the muscle overlying the L3, L4, and L5 DRG was removed. Using fine-tipped rongeurs, the bone surrounding the L4 DRG was carefully removed in a caudal-rostral direction. Bleeding was prevented using gelfoam (Spongostan™; Ferrosan, Denmark). The DRG was washed and kept moist using 0.9 % sterile saline. The position of the mouse was varied between prone and lateral recumbent to orient the DRG in a more horizontal plane. The exposure was then stabilized at the neighbouring vertebrae using spinal clamps (Precision Systems and Instrumentation, VA, USA) attached to a custom-made imaging stage. Finally, the DRG was covered with silicone elastomer (World Precision Instruments, Ltd) to maintain a physiological environment. Prior to imaging, we administered a subcutaneous injection of 0.25 ml 0.9 % sterile saline to keep the mouse hydrated. It was then placed under an Eclipse Ni-E FN upright confocal/multiphoton microscope (Nikon), and the microscope stage was diagonally orientated to optimise focus on the L4 DRG. The ambient temperature during imaging was kept at 32°C throughout. All images were acquired using a 10× dry objective. A 488-nm Argon ion laser line was used to excite GCaMP6s, and the signal was collected at 500–550 nm. Time lapse recordings were taken with an in-plane resolution of 512 × 512 pixels and a partially (∼3/4) open pinhole for confocal image acquisition. All recordings were acquired at 3.65 Hz. Mice were culled with an overdose of sodium pentobarbital at the end of each experiment.

#### photon imaging

Imaging was carried out as previously described. Briefly, adult MrgprD-Cre mice (after viral injection) were deeply anesthetized intraperitoneally with 100 mg/kg ketamine, 15 mg/kg xylazine, and 2.5 mg/kg acepromazine (A7111, Sigma-Aldrich) ^21,22^. Laminectomy was performed to expose L4 DRG, and the spinal columns flanking the laminectomy exposure were clamped with two clamps of a home-made spinal stabilization device to fix the animals. 3% agar solution was used to make a pool for holding Ringer solution (in mM: 126 NaCl, 2.5 KCl, 2 CaCl2, 2 MgCl2, 10 D-Glucose, 10 HEPES, pH = 7.0). Dextran Texas Red (70 kDa, Neutral, D1830, Invitrogen; 1% in saline) was injected intravenously to label the blood vessels, which provide structural landmarks for image registration. The animals were heated with a heating pad during the surgery and imaging to keep the body temperature at 37℃. Warmed Ringer solution was dropped on the exposed spinal cord and DRGs to keep the moisture, and repeatedly changed during the imaging experiment. Animals with the whole spinal stabilization device were fixed under a home-made video-rate 2-photon microscope. A tunable InSight X3 femto-second laser (Spectra-Physics) was set to 940 nm for GCaMP6s and Texas Red imaging. Images were acquired at 32 Hz with an Olympus water-immersion 40x objective at resolution of 0.375 um/pixel.

### Calcium imaging data processing

#### photon recordings

Timelapse recordings were concatenated and scaled to 8-bit in Fiji/ImageJ, Version 1.53. The image analysis pipeline Suite2P (https://github.com/MouseLand/suite2p; v 0.9.2) ^23^ was utilised for motion correction, automatic region of interest (ROI) detection and signal extraction. Further analysis was undertaken with a combination of Microsoft Office Excel 2013, Matlab (2018a) and RStudio (Version 4.02). A region of background was selected, and its signal subtracted from each ROI. To generate normalised ΔF/F0 data, the ROMANO toolbox ProcessFluorescentTraces() function was utilised^24^. This function uses the calculation: ΔF/F0 = (Ft-F0)/F0, where Ft is the fluorescence at time t and F0 is the fluorescence average over a baseline period. ΔF/F0 is expressed as a percentage.

#### photon recordings

The recorded images were processed and analysed as previously reported^19^. Briefly, the RAW image sequences were first converted into TIFF format in ImageJ. Then rigid body translation alignment based on the 2D cross-correlation was performed with a custom-built MATLAB (MathWorks) function to correct for movement (image registration). A rectangular region of interest (ROI) in a region without any visible neuron was drawn as background ROI. The average pixel value inside the background ROI for each frame was subtracted from every pixel in the corresponding frame to remove excess noise. Then small rectangular ROIs were placed manually in the cytoplasm of individual visible neurons. The average fluorescence intensity of a given ROI, Ft, was measured by averaging pixel values inside the ROI. Calcium traces were calculated as, ΔF/F0 = (Ft-F0)/F0, where F0 is the fluorescence value at baseline, which was measured as the average of the first two seconds of Ft. To avoid aberrant amplification due to small F0 values in some neurons (e.g., low basal fluorescence), when it was <1, F0 in the denominator, but not in the numerator, was set to 1. The resulting Ca2+ time series extracted from the image sequences were synchronized with thermal or mechanical stimulus data series. Processing was performed using custom functions written in MATLAB.

An integrated interface within a custom tool written in Spike2 (CED) was used to automatically detect and measure positive responses to stimuli. Raw Ca2+ traces were first smoothed with a 1 s temporal window. Baseline was selected from a period between 1 s after the beginning of the recording and 1 s before the stimulus onset (usually 3 to 5 s in duration). Then, Fb and Fb-max were calculated as average and maximum ΔF/F0 values during baseline, respectively. A response was considered positive when the peak of the Ca2+ trace during stimulation was above Fb+(Fb-max-Fb)*x, where x was a value between 2 and 3, depending on baseline stability, to provide the most reliable detection. Given the relatively slow decay of GCaMP6s, responses with very brief duration (< 0.5 s) were excluded. All traces were also visually inspected to ensure no false positive were included and no false negative missed.

Prior to running these data through the spontaneous activity detection algorithm, it was down-sampled from 32Hz to 3.65Hz (acquisition speed using for training algorithm) using a linear interpolation function in python.

### Mechanical and heat stimulation of the peripheral terminals

Neurons in the leg and foot were stimulated via mechanical stimulation. Specifically, mechanically sensitive afferents were identified by both pinching, using a pair of serrated forceps, brushing with a paintbrush and by moving the foot.

A feedback-controlled Peltier device and a 1 cm × 1 cm thermal probe (TSA-II-NeuroSensory Analyzer, Medoc) were used to deliver noxious heat stimulation in a fast ramp and hold mode to the plantar side of the hind paw. The speed of temperature increase was set at 8°C/s, and temperature decrease was set at 4°C/s. The baseline temperature was set at 25°C, and the duration of the steady state phase of the 50°C stimulation was 5 s.

### Electrical stimulation of the sciatic nerve

In some experiments, the sciatic nerve was dissected at the level of the mid-thigh and freed of any surrounding connective tissue. A custom-made electrical stimulation cuff made of steel wire was fitted under the nerve, and square wave pulses of 1 ms width 5 mA amplitude (suprathreshold) were applied at 0.2 Hz for 5 minutes.

### *In vivo* lidocaine application

Lidocaine hydrochloride (Sigma, UK; dissolved in 0.9 % w/v sodium chloride to a concentration of 74 mM, 2 % w/v) was used to block activity in spontaneously active neurons, thereby revealing their true baseline level of calcium. Lidocaine injection directly into the native sciatic notch during the recording was challenging. Therefore, to increase the accuracy of our injections onto the sciatic nerve, we ‘marked up’ the position of the notch by making a small incision in the skin and surrounding muscle at the level of the midthigh prior to recording. For mice that were used for spontaneous activity testing, no incisions were made in the leg prior to recording baseline activity. To accurately inject onto the nerve, the mouse was removed from the microscope stage, and the sciatic nerve was exposed at the level of the midthigh. The mouse was then returned to the stage, and the imaging plane was carefully realigned. After two minutes of recording, lidocaine was applied to the nerve.

### Machine learning algorithm

The machine learning algorithms used in this study are available as part of the Alan Turing institute Sktime project (https://www.sktime.org/en/stable/)^25,26^. Preliminary investigations demonstrated that two interval-based algorithms were most effective with this dataset: Time Series Forest classifier (TSF) and Random Interval Spectral Ensemble (RISE). Both algorithms are forest-based, allocating intervals of the input time series to individual tree classifiers. The outputs of individual trees are fed forward towards an ensemble which calculates the final model prediction. TSF divides the entire time series into *n* distinct intervals, calculates the mean, standard deviation and slope of each interval and uses these three features for its tree classifiers. In contrast, RISE converts random time intervals to spectral coefficients which are used as features for its tree classifiers.

Model training was performed using an 80:20 ratio train test split in concert with K-fold cross validation (k=5). Model training and serving are available through a Google Collab notebook: https://github.com/sonialouise/ts_class/blob/main/notebooks/SpontaneousActivity.ipynb and all code is available via Github: https://github.com/sonialouise/ts_class.

### Algorithm training and testing

The algorithms were trained on neurons that were deemed ‘active’ or ‘inactive’, and this was determined as follows. Spontaneous activity was induced using the CFA model, with mice imaged on day one or two post-injection. Spontaneously active neurons were differentiated from those which were silent using lidocaine, which, when applied to the nerve in between the inflamed site and the DRG, blocks any incoming action potentials that are being generated in the inflamed paw (**Fig 1 A**). Block by lidocaine was determined by examining calcium traces by eye (assessed by an experienced observer). Fluorescent traces were taken from spontaneously active neurons during the first (pre-lidocaine) and last (5 mins post-lidocaine) 1100 frames (∼5 mins), to generate ‘active’ and ‘inactive’ training data sets, respectively. Traces in which activity pre- and post-lidocaine was ambiguously altered were excluded from training data sets i.e., they may have been active spontaneously, but their activity might have been generated at a site remote from the inflamed paw.

**Figure 1.**
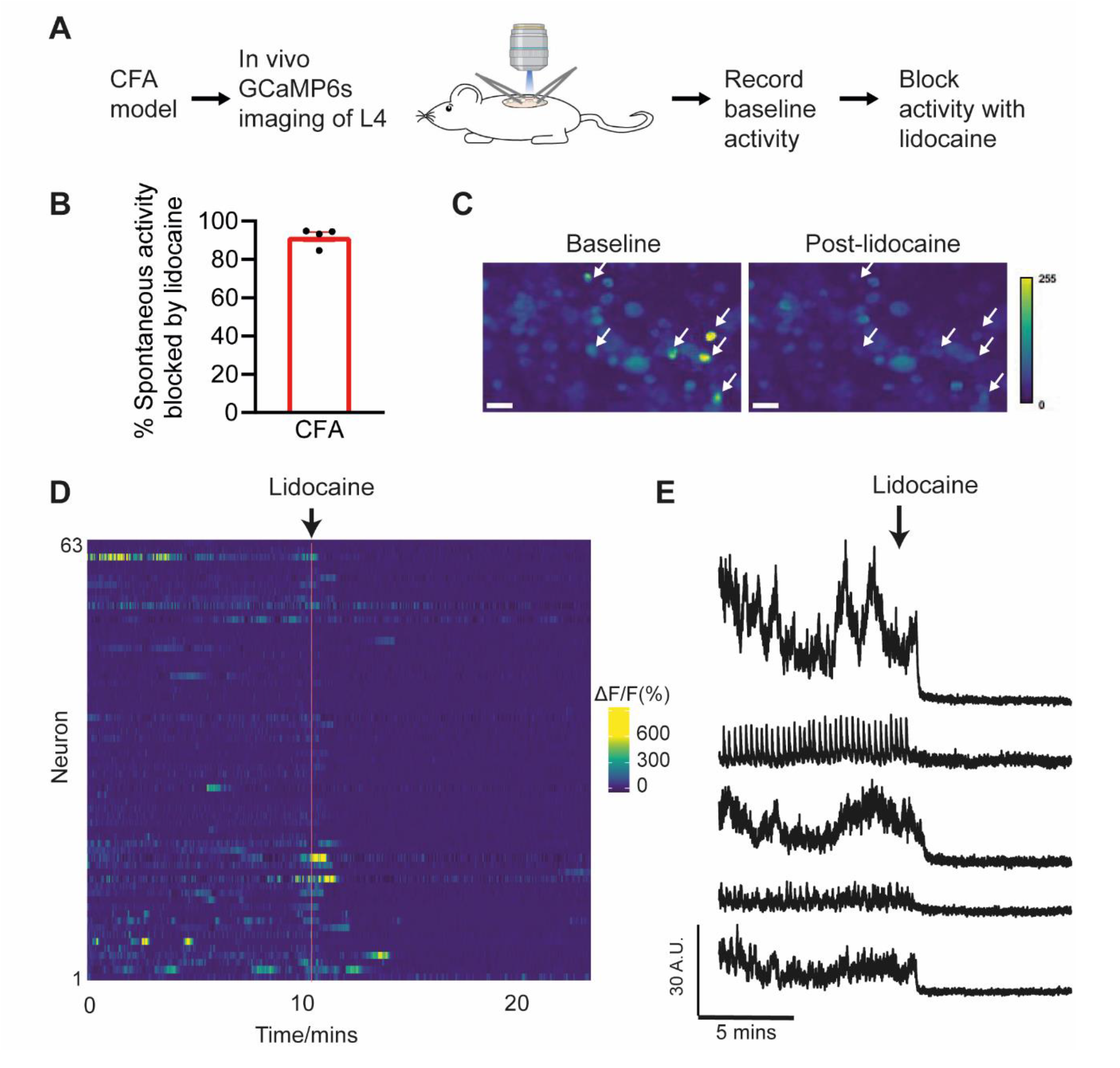
Examination of inflammation-induced spontaneously active calcium transients. Experimental schematic is shown in (A). The bar graph shows the proportion of CFA-induced spontaneously active neurons that were blocked by lidocaine application to the sciatic nerve (B). Example images before and after lidocaine application (C). Images were generated using a z-projection of the standard deviation of 500 frames at baseline (left panel) and post-lidocaine application (right panel). Arrows indicate examples of neurons that were spontaneously active at baseline and silent post-lidocaine. Examples of spontaneously active neurons are shown in the heat map of ΔF/F0 normalised data (D) and in the raw background-subtracted fluorescent traces (E). Lidocaine application to the sciatic nerve (timing indicated by arrows in D & E) in between the inflamed site and the DRG recording site confirmed that spontaneous activity was peripherally generated and revealed the true baseline of these neurons. Scale bar = 50 um. See also Supplementary Video 1.

Electrophysiological experiments indicate that most neurons firing spontaneously in the CFA pain model are nociceptors^27,28^. To ensure that the algorithm was trained on baseline calcium traces in other neuron types, i.e. non-nociceptors, the post-lidocaine data (>5 mins after application) from neurons in the DRG that were not spontaneously active was included in the ‘inactive’ neuron training set. It was assumed that these neurons would be silent because, under normal conditions, activity in sensory neuron is typically generated at the peripheral terminals (which were blocked), and indeed this was true for most experiments (confirmed by visually inspecting traces).

We compared the consequences of training the algorithm with two different types of calcium imaging data: raw fluorescent calcium traces or normalized traces using ΔF/F0. Each version was then tested for prediction accuracy using independent data sets. Unless otherwise stated, sensitivity, specificity and accuracy were calculated as follows:

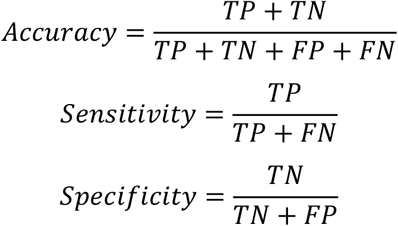

True positive (TP) = the number of neurons with spontaneous activity (as determined by lidocaine). False positive (FP) = the number of neurons incorrectly identified as active. True negative (TN) = the number of neurons without spontaneous activity (as determined by lidocaine). False negative (FN) = the number of neurons incorrectly identified as inactive.

The experiments conducted by our collaborator did not involve application of lidocaine, nor were they conducted in a disease context in which one would expect spontaneous activity. However, they were designed to induce neuronal firing in response to heat and mechanical stimulation. We therefore conducted a blinded trial to assess our algorithm’s ability to detect evoked activity in these 2-photon recordings. Active neurons were determined using a custom written tool in Spike2, which can accurately detect evoked but not spontaneous activity, and then the results were compared to those generated using the machine learning algorithm.

### Quantification and Statistical analysis

Graphing and statistical analysis was undertaken with a combination of Microsoft Office Excel 2013, R Studio (Version 4.02) and GraphPad Prism (Version 8). Details of statistical tests and sample sizes are recorded in the appropriate figure legends. All data plotted represent mean ± SEM. Unpaired student t-tests were used to compare the proportion of spontaneously active neurons between treatment and control groups.

## Results

### Assessment of spontaneous activity in an inflammatory pain model

We chose the CFA model to study spontaneous activity, because a high proportion of sensory neurons are reported spontaneously active at early time points in electrophysiology studies^27,28^. As expected, GCaMP6s mice that were imaged using a 1-photon confocal microscope one to two days post-CFA injection showed increased spontaneous calcium transients that resembled spontaneous activity (Fig 1B & C). Application of lidocaine in between the inflamed hind paw and the DRG recording site blocked the majority of spontaneous calcium transients that were observed in CFA animals (91.9% +/-2.4, n=4 mice). This confirmed that the activity was generated by action potentials, i.e. not caused by movement artefacts, and that it was generated in the periphery (Fig 1B-D, Supplementary Video 1).

Spontaneous activity recorded in the DRG may itself originate in the soma of primary afferents, rather than the axon^29^. Therefore, it should be noted that we were likely unable to inhibit all spontaneous activity through application of lidocaine onto the nerve. Lidocaine application to the DRG was not possible due to our recording set up (see methods), and as a result we were not able to quantify the total number of spontaneously active neurons in the DRG. Indeed, the skill required to apply lidocaine directly onto the nerve while visualising the DRG or, more difficult still, applying lidocaine directly to the DRG (to remove remaining ectopic spontaneous activity), presents a significant barrier to the study of spontaneous activity in the peripheral nervous system. Additionally, the added step of lidocaine application to any parts of the peripheral nervous system often does not fit easily within a given experimental paradigm. We therefore set out to develop an algorithm that could be used on any baseline recordings to accurately predict when a neuron is firing, including on data already recorded for other purposes.

### Development of an algorithm to detect spontaneous activity in *in vivo* calcium imaging recordings

In order to assemble a ‘ground truth’ dataset of spontaneously active neurons, we performed imaging experiments on n=8 CFA mice. In each experiment, lidocaine nerve block was used to distinguish spontaneous activity from baseline noise (Fig 2A). We thought that perhaps a measure of variance, such as SD, could predict spontaneous activity. However, we found that the standard deviation of spontaneously active and inactive segments overlapped more than previously anticipated: 77.9 % of neurons with no activity and 23.1 % of those with activity had SDs of below 15 % (ΔF/F0; Supplementary Fig 1A). We applied basic formulas utilising the SD in order to distinguish activity from noise and could not identify an acceptable compromise between sensitivity and specificity, i.e. increasing the ability of the algorithm to detect true spontaneously active neurons would lead to a very high rate of false positives and vice versa. A threshold for spontaneous activity of >2.5*SD on more than 30 occasions in 5 minutes performed the best on our ground truth data, but prediction accuracies were not reproducible when testing on independent CFA experiments (Accuracy = 78.0% +/-0.6; Supplementary Fig 1B). We therefore decided to investigate whether supervised machine learning algorithms specialised for time series data could provide better predictive accuracies. We trained two different algorithms, using the visual lidocaine block as ‘ground truth’ data, on both ΔF/F0 normalised and non-normalised raw fluorescent data (1308 neurons, 272 active and 1036 inactive from n=8 CFA mice). The two algorithms that were tested were the Time Series Forest classifier (TSF) and Random Interval Spectral Ensemble RISE). Both algorithms are forest-based and were selected because they performed best on our datasets in preliminary testing.

**Figure 2.**
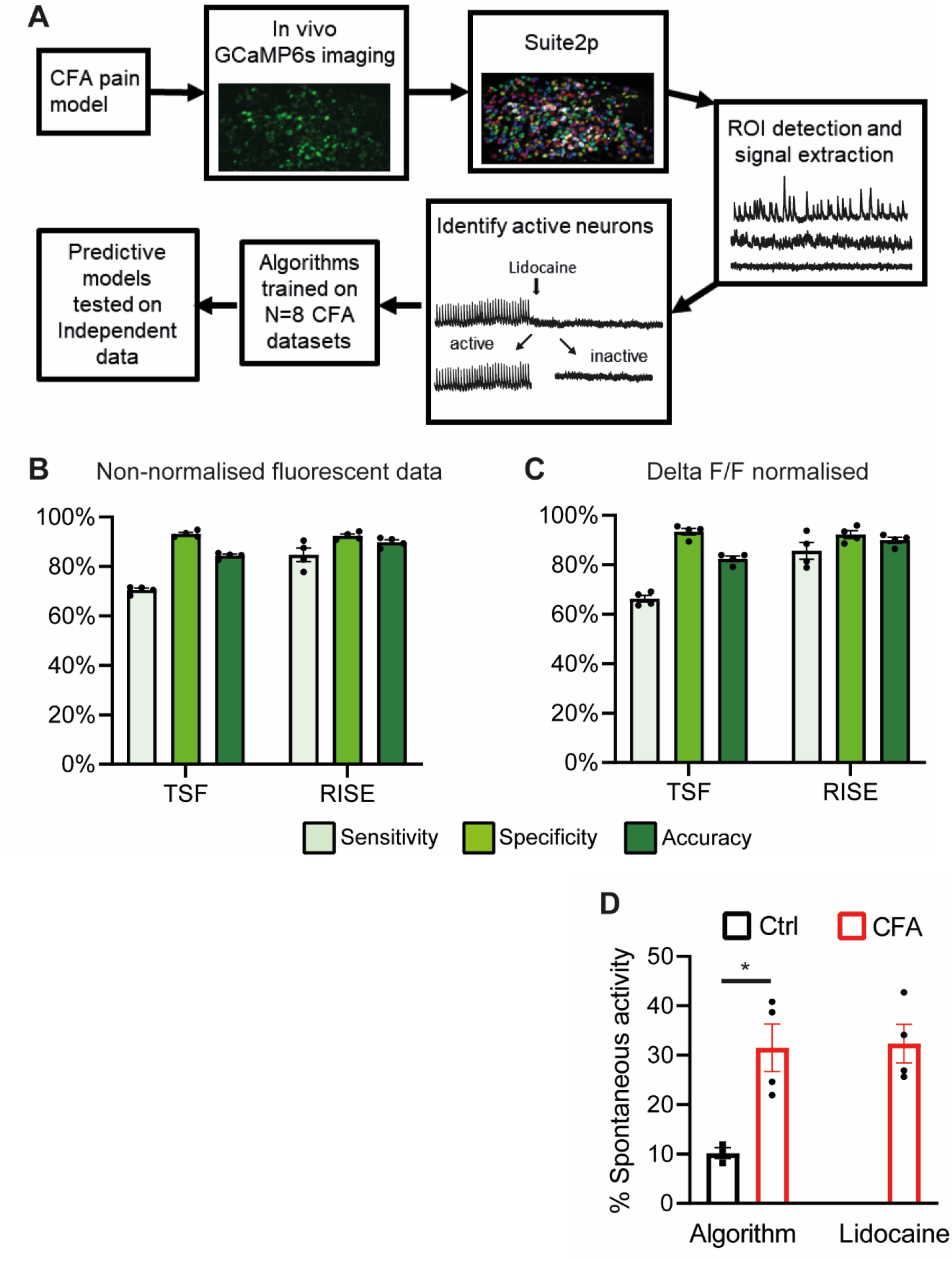
Training machine learning algorithms to detect spontaneous activity in GCaMP6s DRG recordings. Schematic shows the workflow of how CFA induced spontaneously active neurons were identified for algorithm training (A). The algorithms were tested on n=4 independent CFA experiments, after having been trained on either non-normalised (B) or ΔF/F0 normalised (C) data. Sensitivity = the ability of the algorithm to correctly identify when a neuron is spontaneously active. Specificity = the ability of the algorithm to correctly identify when a neuron is not active. Accuracy = the ability of the algorithm to differentiate spontaneously active from inactive neurons. Lidocaine was used to establish ground truth data, i.e. determine whether a neuron was spontaneously active or not. TSF – time series forest classifier. RISE – random interval spectral ensemble. The proportion of neurons in the L4 DRG that were predicted to have spontaneous activity in control and CFA animals on day 1 is shown in (D). Left side – algorithm prediction, right side – conventional quantification by experimenter. Note that our datasets only include information on spontaneously active neurons blocked by lidocaine application to the nerve i.e. not those that might have had activity originating in the cell soma. n=3/4 group. Bar graphs represent mean ± SEM.

Upon testing on n=4 independent CFA datasets (225-316 neurons/ experiment) we found that the normalisation method (raw fluorescence vs deltaF/F0) had little impact on the predictive accuracy of our algorithms (Fig 2 B and C). However, for both normalisation methods, the RISE algorithm predicted spontaneous activity more accurately than the TSF algorithm (TSF accuracy = 84.4 % +/-0.6, RISE accuracy 87.2 % = +/-1.0; TSF accuracy = 82.4 % +/-1.1, RISE accuracy = 90.0 % +/-1.2, respectively for non-normalised and normalised data; Fig 1C). Going forward, we therefore opted to use the RISE algorithm trained on ΔF/F0 normalised data, as it performed best in these initial tests, and is likely more generalisable (ΔF/F0 being the most dominant processing step in the field).

Reducing the number of frames the algorithm was trained and tested on from 1100 to 550 or 200 did not markedly reduce the algorithms prediction accuracy performance (RISE accuracies = 88.9% +/-1.7, 88.3% +/-1.7 for 550 and 200 frame training respectively; n=4 mice; Supplementary Fig 2A & B). When the algorithms trained on GCaMP6s normalised data were tested on CFA mice imaged with GCaMP8s, the prediction accuracy was somewhat reduced (RISE accuracy = 86.0% +/-2.8, n=2; Supplementary Fig 3A). Surprisingly, given that GCaMP8s is reported to be a more sensitive calcium indicator than GCaMP6s^30^ (Supplementary Fig 3B), this was caused by a reduction in sensitivity, i.e. the ability of the algorithm to correctly identify spontaneously active neurons (70.3% +/-4.4, n=2).

Finally, the RISE algorithm trained on ΔF/F0 data predicted 31.5% (+/-4.8) of neurons to be active 1 day following CFA injection; n=4 independent mice), which was significantly greater (p=0.014; unpaired t-test) than the number of spontaneously active neurons observed in control mice (10.2% +/-1.1; n=3; Fig 2D-left). This was very similar to the number of neurons found to be spontaneously active by an experimenter, using the ‘gold standard’ lidocaine method of identification (32.3% +/-3.9; Fig 2D – right).

### Testing the robustness of the algorithm using additional datasets and disease models

Next, we tested the RISE algorithm’s ability to predict when a neuron was firing spontaneously in an antigen-induced arthritis (AIA) model of joint pain (Fig 3A). After immunisation, mice that underwent intraarticular injections of mBSA showed an increase in the size of the ipsilateral joint compared to its contralateral, uninjected counterpart (ipsi/contra ratio = 1.31 +/-0.07; n=6). This was significantly different (p=0.011, unpaired t-test) to the saline injected group (ipsi/contra ratio = 1.04 +/-0.01; n=4; Fig 3B). Consistent with the results in the CFA model, the RISE algorithm accurately detected spontaneous activity in both treatment groups (accuracies = 85.9 % +/-2.1 and 94.4 % +/-0.5 in mBSA and control groups, respectively).

**Figure 3.**
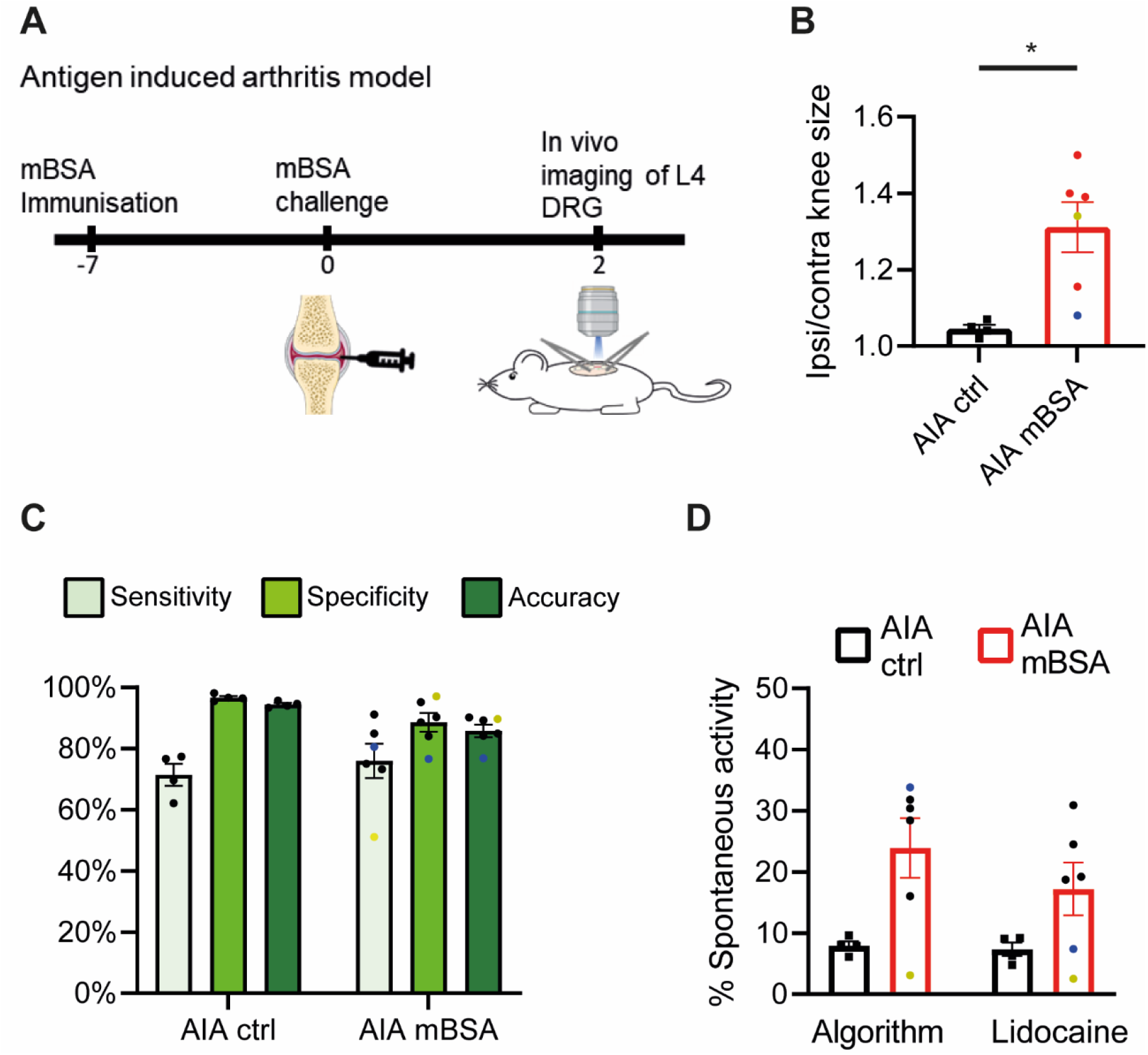
Assessment of spontaneous activity in an arthritis model. Experimental schematic illustrating how the AIA model is induced (A). The ratio of ipsilateral to contralateral knee size in mice 2 days after receiving an intraarticular injection of saline (ctrl) and mBSA (B). The results of the RISE algorithm’s ability to detect spontaneous activity in the AIA model (C). The proportion of neurons in the L4 DRG with spontaneously activity in control and AIA animals on day 2 is shown in (D). RISE algorithm prediction is shown on the left, while conventional quantification by an experimenter using lidocaine-induce silencing is on the right. The yellow dots denote an experiment in which the GCaMP6s labelling efficiency was unusually low, i.e. the laser power was double that of what is typical; consequently, the sensitivity was reduced in this experiment. The blue dots denote an experiment that had a bad movement artefact; consequently the specificity was reduced in this experiment. n=4/group. ΔF/F0 normalised CFA data were used for training. Bar graphs represent mean ± SEM.

The yellow dot represents an experiment where the GCaMP6s labelling was suboptimal, i.e the laser strength required to visualise neurons was doubled compared to our standard settings (10 % vs 5 %). Although the knee was clearly inflamed in this animal (ispi/contral ratio = 1.33), very few neurons were spontaneously active (2.5% as determined by lidocaine), and hence sensitivity of the RISE algorithm was reduced (50%). Nevertheless, thanks to its continuing high specificity, RISE appeared relatively robust to such a variation in labelling with overall accuracy levels only slightly lower than average (89.7%). In contrast, when specificity was reduced to 76.6%, due to a particularly bad movement artefact, the accuracy of the algorithm was more noticeably affected (accuracy = 76.9%; represented by blue dots in Fig 3C).

Using the RISE algorithm, spontaneous activity was identified in 23.9% (+/-4.9; n=6) of neurons following mBSA injection, which was similar to the percentage that identified by an experimenter using lidocaine (17.2% +/-4.3; Fig 3D). There was a non-significant increase in the number of neurons identified with spontaneous activity in the mBSA group compared to the saline injected control group (7.9 % +/-0.7 active; n=4; using RISE algorithm predictions, p=0.29, unpaired t-test; Fig 3D).

To determine if the algorithm could detect low frequency firing that was just above the level of background noise, the sciatic nerve was stimulated at suprathreshold strength at 0.05-0.2 Hz (Fig 4 A – left panel and see Supplementary Video 2). The RISE algorithm was able to detect low frequency electrical activity 80.9 % of the time (+/-4.8, n=4; Fig 4 B). In contrast, very few neurons were predicted active when all activity was blocked by lidocaine (6.6 % +/-1.4, n=4; Fig 4 C).

**Figure 4.**
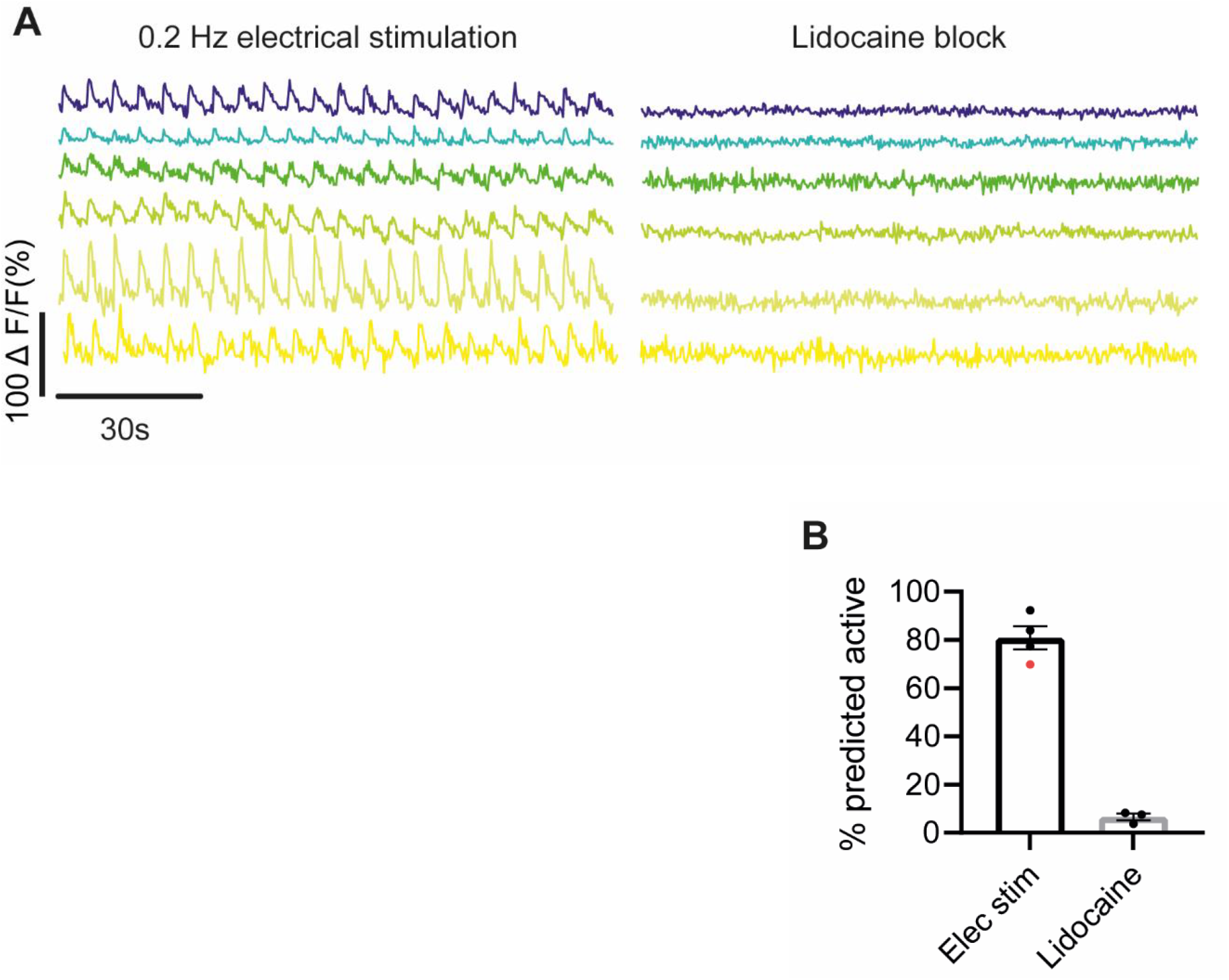
The RISE algorithm can detect the majority of neurons firing at low frequencies. Example ΔF/F0 normalised data traces of neurons firing at 0.2Hz and after lidocaine block are shown in (A). Bar graph shows the proportion of neurons that were predicted active by the RISE classifier when stimulated at 0.05 - 0.2 Hz vs. following lidocaine block (B). The algorithm could detect low frequency neuronal activity, and very few neurons were predicted active when all neuronal activity was blocked by lidocaine. See also Supplementary Video 2. ΔF/F0 normalised CFA data were used for training. Bar graphs represent mean ± SEM. Red dot denotes experiment in which the nerve was electrically stimulated at 0.05 Hz, all other stimulations were performed at 0.2 Hz.

To ensure our algorithm was robust enough to detect neuronal activity in different experimental settings, we tested it on data collected in a different laboratory under different conditions, i.e. both a different microscope configuration and acquisition rate. These 2-photon data required additional normalisation beforehand. We smoothed them using a moving average and down-sampled the acquisition rate from 32 Hz to 3.65 Hz using a linear interpolation function in Python (Fig 5A). The RISE algorithm could accurately detect activity in 1-minute-long 2-photon microscope recordings (accuracy = 94.0 % +/-2.2; 30-72 neurons/mouse, n=6 mice; Fig 5B), despite being trained on data recorded with a single photon microscope.

**Figure 5.**
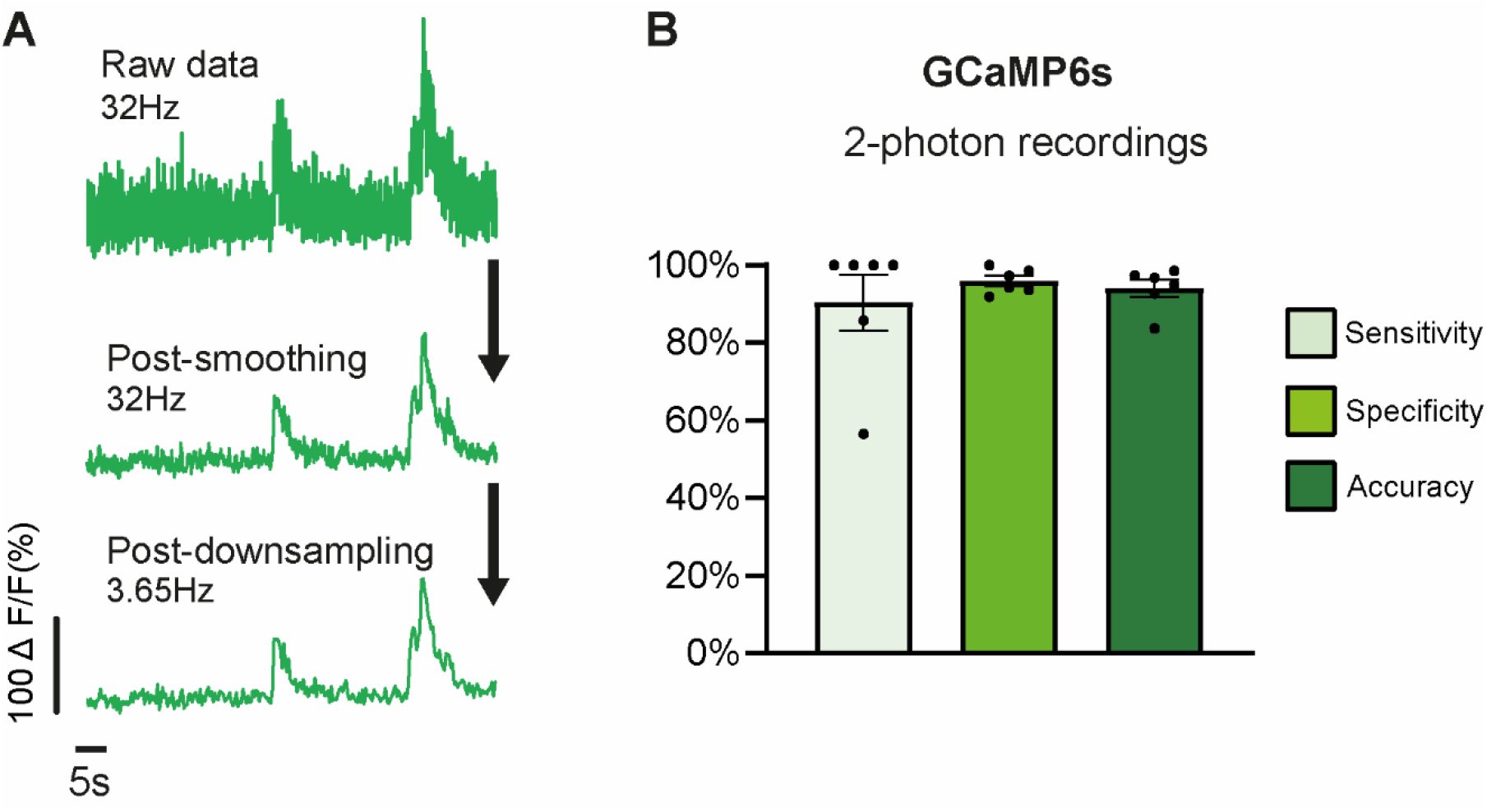
The RISE algorithm can detect GCaMP6s activity collected in a different lab using a different microscope configuration. Pre-processing steps necessary to convert 2-photon GCaMP6s data to a format that could be used for testing (A). Example original ΔF/F0 normalised data traces (top) vs. after smoothing (middle trace) and down-sampling (bottom trace). The results of the RISE algorithm’s ability to detect evoked activity in experiments performed with 2-photon microscope (B). Note that in this case, sensitivity, specificity and accuracy were calculated using experimenter-determined response rates provided by the other lab in a blinded fashion (see methods for further details). Data collected from n=6 mice. ΔF/F0 normalised CFA data were used for training. Bar graphs represent mean ± SEM.

## Discussion

Using traditional electrophysiological techniques to study spontaneous activity in sensory neurons has been challenging. In this study, we set out to determine if *in vivo* calcium imaging can be used as an alternative means to study this type of activity in larger numbers of neurons at once. Indeed, we found that spontaneously active neurons can be visualised in 1-photon recordings using GCaMP indicators in a CFA model of pain. Using these data, we trained two supervised machine learning algorithms to identify whether either could successfully identify when a neuron was spontaneously active. We found that the RISE algorithm, trained on normalised ΔF/F0, performed particularly well. We tested it on independent CFA data and on an arthritis pain model and were able to demonstrate that it is robust enough for detecting inflammation-induced sensory neuron spontaneous activity in both cases. Further, the algorithm can accurately predict activity on data collected by a different laboratory using a 2-photon microscope, suggesting that it is robust enough for detecting both spontaneous and evoked sensory neuron activity across different GCaMP6s datasets.

The proportion of neurons with spontaneous activity in control (∼10 %) and CFA inflamed mice (∼32 % on day 1) is broadly in line with that previously reported in the literature via electrophysiological studies in rats^27,31,32^. So far, few studies have reported on spontaneous activity in sensory neurons using calcium indicators *in vivo*. In comparison to our baseline data, two studies using a GCaMP3 transgenic mouse line reported much lower levels (0.13-0.19 %) of spontaneously active calcium transients^33,34^. This difference is likely because the overall acquisition rate in these recordings was a lot slower in comparison to ours (<0.2 Hz vs 3.65 Hz). At the temporal resolution of 0.2 Hz, lower frequency firing would not have been detectable. A separate study using a GCaMP6s transgenic mouse line also found a lower proportion of neurons that were spontaneously active at baseline in anaesthetised mice (∼4%), which might be because different anaesthetics were used: we used isoflurane in combination with urethane, while the other study used ketamine and xylene. Indeed, anaesthesia can impact peripheral neuron excitability, with the proportion of neurons with spontaneous activity reportedly higher (∼14 %) in awake animals^35^.

There are many advantages of using *in vivo* imaging to study spontaneous activity over electrophysiology. One of the major conundrums of using electrophysiology is that it is difficult to record spontaneously active nociceptive neurons (e.g. compared to A fibres), because they are the smallest in size. In contrast, all sizes and subgroups of sensory neuron can be visualised using GCaMP calcium indicators^17,19^, thereby reducing the bias towards recording from a particular fibre type. Electrophysiology is low throughput – 10s of neurons/experiment, whilst we typically record from ∼400-500 neurons/DRG in an imaging experiment; imaging is therefore not only faster, but the simultaneous recording of many neurons also allows for the examination of coordinated patterns of spontaneous firing. Finally, neurons must be activated to visualise them when using electrophysiology, which can cause neuronal sensitisation and increase spontaneous firing when done repeatedly via receptive field testing^16^. In contrast, no activation is required for visualising neurons prior to starting the recordings using *in vivo* imaging.

Good signal to noise is required to be able to visualise and detect low frequency neuronal firing in *in vivo* imaging experiments with GCaMP6s. Reduced signal to noise can be caused by poor GCaMP labelling. This may result in a reduction of the number of spontaneously active neurons that are capable of being visualised. Indeed, in one of our AIA experiments with particularly bad signal to noise, only 2.5% of neurons were spontaneously active, despite the knee being clearly inflamed (knee size ratio = 1.33). Although the accuracy was not reduced in this experiment, the sensitivity was affected (see yellow dot in Fig 3C), and this is likely because the prevalence of spontaneous activity (i.e. the number of true positives) was so low that the number of false negatives has a greater impact on the calculation. Movement artefacts, e.g. from breathing, can also impact signal to noise. Such artefacts can normally be mitigated by ensuring the spinal vertebra are clamped adequately, so that the preparation is stable, and by using image registration algorithms. However, sometimes movement artefacts are difficult to completely remove, and this can lead to a reduction in the accuracy of detecting spontaneous activity using our trained machine learning algorithm (see example indicated by blue dots in Fig 3).

We used the normalised ΔF/F0 data for training our algorithms, because it produced accuracy results that were almost identical to that of the non-normalised data. One might find this somewhat surprising, given the baseline that we are normalising against should already be raised in a spontaneously active neuron, thus decreasing the overall amplitude of any signal. However, this turned out not to be the case when examining inflammation-induced spontaneous activity. It is likely that this is because spontaneously active neurons fire at relatively low rates (<1 Hz) during inflammation^27,36^, too low to cause saturation of GCaMP6s^17^. Indeed, it seems unlikely that the algorithms trained on ΔF/F0 normalised data would be able to accurately predict spontaneously active neurons firing at higher frequencies (>5 Hz), which is a caveat of using our currently trained versions. However, as there are now faster and more sensitive GCaMPs, which are reported to be able to follow firing frequencies of ≥10 Hz^30^, it seems likely that this obstacle can be overcome by retraining the algorithms on novel data collected using these indicators.

Indeed, in the future, we aim to improve the robustness of our algorithm by retraining with data generated with different indicators. This will be necessary, as we have found that with its current training set based on GCaMP6s data, the RISE algorithm cannot detect activity as accurately when it is derived from neurons containing GCaMP8s as an indicator. This is somewhat surprising, as GCaMP8s is known to be more sensitive than GCaMP6s, but probably due to differences in kinetics: consistent with previous reports^30^, we found the half-life decay time of GCaMP6s to be slower than that of GCaMP8s (see Supplementary Fig 3B).

There are often difficulties setting up a new toolbox in python for inexperienced users. To make our algorithm easy for others to use, we have generated a cloud-based version that is available as a Google Colaboratory Notebook. Input data can be acquired at any frame rate, but must be below 5 minutes in duration. For more accurate predictions, we recommend inputting data that has a duration as close to 5 minutes as possible. Both spontaneous and evoked GCaMP6s activity should be detectable with our current algorithm, as long as signal-to-noise ratios are not low i.e. by minimising movement artefacts and ensuring indicator labelling is not weak. Users can also use the Notebook to retrain the algorithm based on their own training sets – however this will require them generating their own ‘ground-truth’ activity data using lidocaine.

## Conclusion

We conclude that *in vivo* imaging with GCaMP indicators is a suitable technique for detecting spontaneous activity in models of pain. We have overcome the difficulty of differentiating spontaneous activity from noise by utilising a machine learning algorithm, which, when trained, can accurately detect GCaMP6s activity in both 1-photon and 2-photon recordings. Our trained algorithm will provide a useful tool for those performing *in vivo* imaging experiments in the pain and wider-neuroscience community.

## Acknowledgments

FD is the recipient of a Medical Research Foundation Prize (MRF-160-0015-ELP-DENK-C0844). FD & GG are funded by an *Advanced Pain Discovery Platform UKRI MRC grant* (MR/W027518/1). KIC is funded by the University of Nottingham Anne McLaren Fellowship. We would like to thank and commemorate our former mentor Stephen McMahon, who was involved in the early stages of this project, but sadly passed away in late 2021.

## Author contributions

G.G. conceived, designed, and performed experiments, analysed data, and wrote the manuscript; S.I conceived experiments, wrote code for training machine learning algorithms and Google Collab notebook, participated in writing paper, provided conceptual input on the project and the manuscript. K.C. wrote code for the algorithms, provided conceptual input, and corrected the manuscript. F.D. provided conceptual input to the manuscript and edited the manuscript. F.W performed experiments and analysed data.

## Data availability

All the training data will be deposited on the GitHub (https://github.com/sonialouise/ts_class). All other data are available upon reasonable request from the corresponding author.

## Code availability

A cloud-based version of the spontaneous activity detection algorithm is available as a Google Colaboratory Notebook: https://github.com/sonialouise/ts_class/blob/main/notebooks/SpontaneousActivity.ipynb

## References

1 Orstavik, K. et al. Pathological C-fibres in patients with a chronic painful condition. Brain 126, 567–578 (2003).

2 Namer, B. et al. Microneurographic assessment of C-fibre function in aged healthy subjects. J Physiol 587, 419–428, doi:10.1113/jphysiol.2008.162941 (2009).

3 Serra, J. et al. Microneurographic identification of spontaneous activity in C-nociceptors in neuropathic pain states in humans and rats. Pain 153, 42–55, doi:S0304-3959(11)00508-2, 10.1016/j.pain.2011.08.015 (2012).

4 Kleggetveit, I. P. et al. High spontaneous activity of C-nociceptors in painful polyneuropathy. Pain 153, 2040–2047, doi:10.1016/j.pain.2012.05.017 (2012).

5 Ochoa, J. L., Campero, M., Serra, J. & Bostock, H. Hyperexcitable polymodal and insensitive nociceptors in painful human neuropathy. Muscle Nerve 32, 459–472, doi:10.1002/mus.20367 (2005).

6 Campero, M., Bostock, H., Baumann, T. K. & Ochoa, J. L. A search for activation of C nociceptors by sympathetic fibers in complex regional pain syndrome. Clinical neurophysiology: official journal of the International Federation of Clinical Neurophysiology 121, 1072–1079, doi:10.1016/j.clinph.2009.12.038 (2010).

7 Serra, J. et al. Hyperexcitable C nociceptors in fibromyalgia. Annals of Neurology 75, 196–208, doi:https://doi.org/10.1002/ana.24065 (2014).

8 Namer, B. et al. Pain relief in a neuropathy patient by lacosamide: Proof of principle of clinical translation from patient-specific iPS cell-derived nociceptors. EBioMedicine 39, 401–408, doi:10.1016/j.ebiom.2018.11.042 (2019).

9 Qu, L. & Caterina, M. J. Enhanced excitability and suppression of A-type K(+) currents in joint sensory neurons in a murine model of antigen-induced arthritis. Scientific reports 6, 28899–28899, doi:10.1038/srep28899 (2016).

10 Devor, M., Wall, P. D. & Catalan, N. Systemic lidocaine silences ectopic neuroma and DRG discharge without blocking nerve conduction. Pain 48, 261–268, doi:0304-3959(92)90067-L (1992).

11 Tal, M. & Eliav, E. Abnormal discharge originates at the site of nerve injury in experimental constriction neuropathy (CCI) in the rat. Pain 64, 511–518 (1996).

12 Goodwin, G. & McMahon, S. B. The physiological function of different voltage-gated sodium channels in pain. Nat Rev Neurosci 22, 263–274, doi:10.1038/s41583-021-00444-w (2021).

13 Jayakar, S. et al. Developing nociceptor-selective treatments for acute and chronic pain. Sci Transl Med 13, eabj9837, doi:10.1126/scitranslmed.abj9837 (2021).

14 Goodwin, G., McMurray, S., Stevens, E. B., Denk, F. & McMahon, S. B. Examination of the contribution of Nav1.7 to axonal propagation in nociceptors. bioRxiv, 2021.2003.2012.435114, doi:10.1101/2021.03.12.435114 (2021).

15 Goodwin, G., Bove, G. M., Dayment, B. & Dilley, A. Characterizing the Mechanical Properties of Ectopic Axonal Receptive Fields in Inflamed Nerves and Following Axonal Transport Disruption. Neuroscience 429, 10–22, doi:10.1016/j.neuroscience.2019.11.042 (2020).

16 Bove, G. M. & Dilley, A. The conundrum of sensitization when recording from nociceptors. J Neurosci Methods 188, 213–218, doi:S0165-0270(10)00087-7, 10.1016/j.jneumeth.2010.02.010 (2010).

17 Chisholm, K. I., Khovanov, N., Lopes, D. M., La Russa, F. & McMahon, S. B. Large Scale In Vivo Recording of Sensory Neuron Activity with GCaMP6. eneuro 5, ENEURO.0417-0417., doi:10.1523/eneuro.0417-17.2018 (2018).

18 Chen, T. W. et al. Ultrasensitive fluorescent proteins for imaging neuronal activity. Nature 499, 295–300, doi:10.1038/nature12354 (2013).

19 Wang, F. et al. Sensory Afferents Use Different Coding Strategies for Heat and Cold. Cell Reports 23, 2001–2013, doi:https://doi.org/10.1016/j.celrep.2018.04.065 (2018).

20 Weiss, M. et al. IRF5 controls both acute and chronic inflammation. Proceedings of the National Academy of Sciences 112, 11001–11006, doi:10.1073/pnas.1506254112 (2015).

21 Vrontou, S., Wong, A. M., Rau, K. K., Koerber, H. R. & Anderson, D. J. Genetic identification of C fibres that detect massage-like stroking of hairy skin in vivo. Nature 493, 669–673, doi:10.1038/nature11810 (2013).

22 Davalos, D. & Akassoglou, K. In vivo imaging of the mouse spinal cord using two-photon microscopy. J Vis Exp, e2760, doi:10.3791/2760 (2012).

23 Pachitariu, M. et al. Suite2p: beyond 10,000 neurons with standard two-photon microscopy. bioRxiv, 061507, doi:10.1101/061507 (2017).

24 Romano, S. A. et al. An integrated calcium imaging processing toolbox for the analysis of neuronal population dynamics. PLOS Computational Biology 13, e1005526, doi:10.1371/journal.pcbi.1005526 (2017).

25 Markus Löning, A. B., Sajaysurya Ganesh, Viktor Kazakov, Jason Lines, Franz Király. A unified interface for machine learning with time series. arXiv preprint arXiv:1909.07872 (2022).

26 Löning, M. et al. sktime/sktime: v0.13.4. doi:https://doi.org/10.5281/zenodo.7117735 (2022).

27 Djouhri, L., Koutsikou, S., Fang, X., McMullan, S. & Lawson, S. N. Spontaneous pain, both neuropathic and inflammatory, is related to frequency of spontaneous firing in intact C-fiber nociceptors. J Neurosci 26, 1281–1292 (2006).

28 Xiao, W. H. & Bennett, G. J. Persistent low-frequency spontaneous discharge in A-fiber and C-fiber primary afferent neurons during an inflammatory pain condition. Anesthesiology 107, 813–821, doi: 10.1097/01.anes.0000286983.33184.9c,00000542-200711000-00019 (2007).

29 Wall, P. D. & Devor, M. Sensory afferent impulses originate from dorsal root ganglia as well as from the periphery in normal and nerve injured rats. Pain 17, 321–339 (1983).

30 Zhang, Y. R., Márton; Bushey, Daniel; Zheng, Jihong; Reep, Daniel; Broussard, Gerard Joey; et al. jGCaMP8 Fast Genetically Encoded Calcium Indicators. Janelia Research Campus. Online resource., doi:https://doi.org/10.25378/janelia.13148243.v1 (2020).

31 Wu, G. et al. Early onset of spontaneous activity in uninjured C-fiber nociceptors after injury to neighboring nerve fibers. J.Neurosci. 21, RC140 (2001).

32 Xiao, W. H. & Bennett, G. J. Chemotherapy-evoked neuropathic pain: Abnormal spontaneous discharge in A-fiber and C-fiber primary afferent neurons and its suppression by acetyl-L-carnitine. Pain 135, 262–270, doi:S0304-3959(07)00303-X;10.1016/j.pain.2007.06.001 (2008).

33 Miller, R. E. et al. Visualization of Peripheral Neuron Sensitization in a Surgical Mouse Model of Osteoarthritis by In Vivo Calcium Imaging. Arthritis Rheumatol 70, 88–97, doi:10.1002/art.40342 (2018).

34 Ishida, H. et al. In Vivo Calcium Imaging Visualizes Incision-Induced Primary Afferent Sensitization and Its Amelioration by Capsaicin Pretreatment. J Neurosci 41, 8494–8507, doi:10.1523/jneurosci.0457-21.2021 (2021).

35 Chen, C. et al. Long-term imaging of dorsal root ganglia in awake behaving mice. Nature Communications 10, 3087, doi:10.1038/s41467-019-11158-0 (2019).

36 Satkeviciute, I., Goodwin, G., Bove, G. M. & Dilley, A. The time course of ongoing activity during neuritis and following axonal transport disruption. J Neurophysiol, doi:10.1152/jn.00882.2017 (2018).

