## Supplementary data for "Assessing spontaneous sensory neuron activity using *in vivo* calcium imaging"

### Supplementary Figures

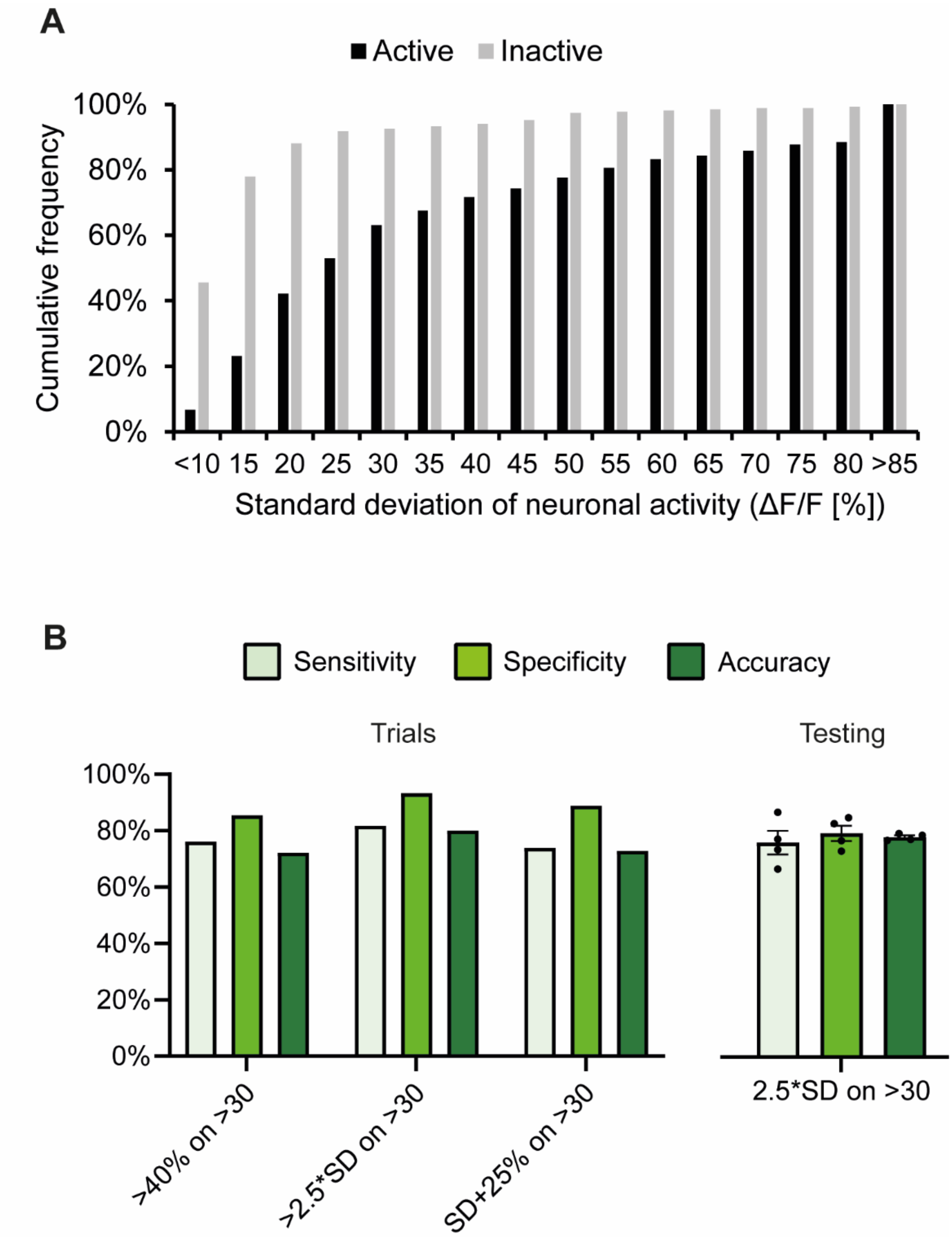


**Supplementary Figure 1. Thresholding data based on standard deviation for predicting spontaneous activity**. *The cumulative frequency histogram (A) shows that the standard deviation of ΔF/F0 normalised spontaneously active (active) and silent (inactive) calcium traces overlap and thus cannot be utilised alone to predict spontaneous activity (n=268 neurons in each group). Note that the same neurons were used in each group, i.e. the activity period was pre-lidocaine and the inactive period was post-lidocaine block. Various formulas were trialled to try and distinguish spontaneous activity from baseline noise (B). A threshold of a rise in calcium 2.5 times the standard deviation on more than 30 occasions in 5 minutes performed the best on the training data set (B-left panel; 268 active/inactive neurons from n=8 CFA mice), but accuracy was reduced when testing on independent data (n=4 CFA mice; B-right panel). Data in histogram is binned into 5% intervals.*

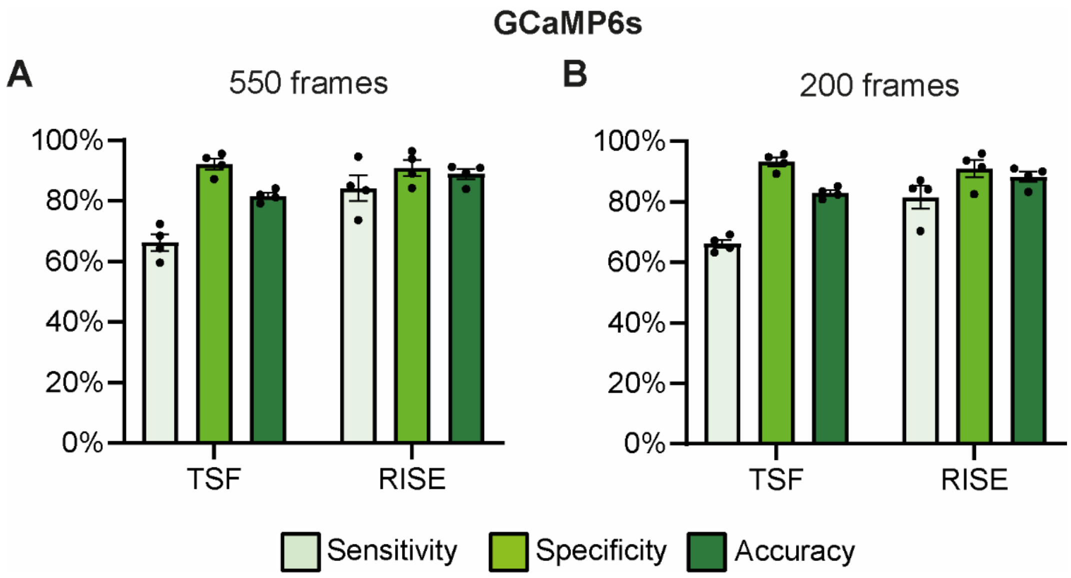


**Supplementary Figure 2. Varying the number of frames used for training did not markedly reduce prediction accuracy of the RISE algorithm**. *Graphs show the results of testing the algorithm's ability to detect spontaneous activity when the frame length used for training was 550 (~ 2.5 minutes; A) and 200 (~ 1 minute; B). Bar graphs represent mean ± SEM.*

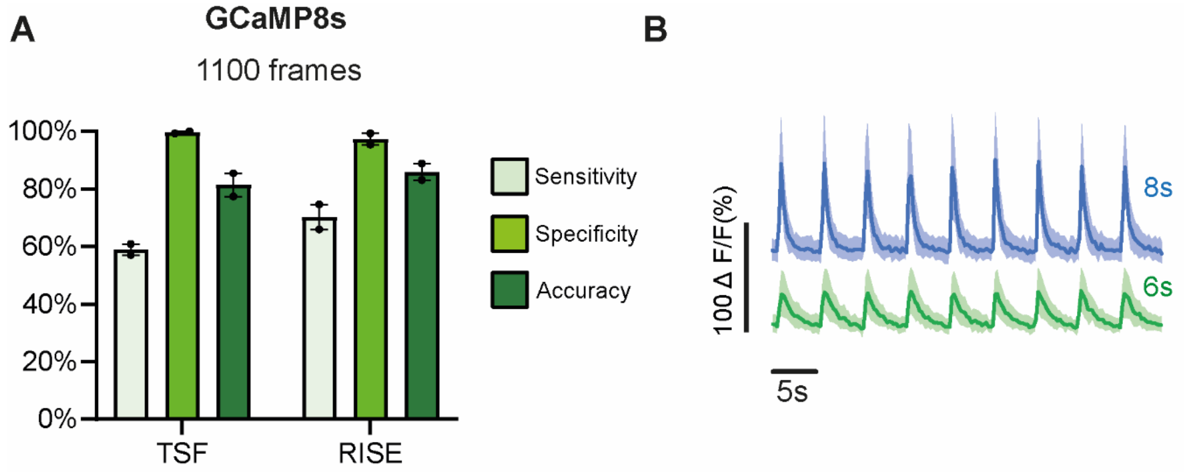


**Supplementary Figure 3. Testing results on CFA data collected with GCaMP8s.** *The graph in (A) shows the results of testing the algorithm's ability to detect spontaneous activity when the calcium indicator used was GCaMP8s. Note that ΔF/F0 normalised CFA data collected using GCaMP6s was used for training. (B) plots averaged ΔF/F0 normalised traces of neurons firing in response to 0.2 Hz electrical stimulation for GCaMP8s (top) and GCaMP6s (bottom). Note that the decay time was slower for GCaMP6s. n=118 from 1x GCaMP8s mouse, n=78 neurons from 1x GCaMP6s mouse. Bar graphs and traces represent mean ± SEM.*

**Links to supplementary videos:**

The videos can be downloaded from the following links. Open the link and use the small menu (button with three small dots) on the right hand side to access the download button.

**Video 1. Inflammation induced spontaneous activity is blocked by peripheral lidocaine application**

https://osf.io/q6u49

*Example In vivo time lapse recording of L4 DRG cell bodies labelled with GCaMP6s. Recording show the left L4 DRG 1-day post-injection of CFA into the left hind paw pre- and post-lidocaine application to the sciatic nerve. Timescale = minutes: seconds*

**Video 2. Example neurons during electrical stimulation and after lidocaine block**

https://osf.io/9qahm

*Example In vivo time lapse recording of L4 DRG cell bodies labelled with GCaMP6s. Recordings show neuronal responses following 0.2Hz electrical stimulation of the sciatic nerve, and after 5-mins lidocaine application to the sciatic nerve. Timescale = minutes: seconds*
